# Sperm-mediated regulation of region-specific responses in the oviduct during establishment of pregnancy in mice

**DOI:** 10.1101/2022.04.18.488702

**Authors:** Ryan M. Finnerty, Daniel J. Carulli, Wipawee Winuthayanon

**Author notes:** Corresponding author: Address: 1770 Stadium Way, Pullman, WA 99164.

## Abstract

The oviduct comprises 4 main regions: infundibulum (oocyte pick-up), ampulla (fertilization), isthmus (sperm capacitation and reservoir, preimplantation embryonic development), and uterotubal junction (UTJ; sperm and embryo transport). Mounting evidence in livestock and rodents suggest that gametes alter gene expression in secretory and ciliated epithelial cells of the oviduct. To elucidate whether adaptive interactions between the oviduct and gamete/embryo exist, we performed bulk RNA-sequencing on oviductal tissues collected from infundibulum+ampulla (IA) or isthmus+UTJ (IU) at various developmental stages (0.5, 1.5, 2.5-, and 3.5-days post coitus (dpc)) in mice. Samples were also collected during days 0.5, 1.5, 2.5, and 3.5 of pseudopregnancy (dpp). We found a strong region (IA vs. IU)-specific expression of large clusters of genes. The transition from 0.5 dpc to other pregnancy timepoints induces large sets of differentially expressed genes (DEGs) in pregnancy and pseudopregnancy in both IA and IU regions. Specifically, genes involved in pro-inflammatory responses were detected in both IU and IA regions. The presence of sperm at 0.5 dpc induces DEGs involved in pro-inflammatory responses in the IU region with an enrichment of biological processes for inflammatory cytokines, macrophage, and neutrophil recruitment. Additionally, DEGs are enriched in mitogen-activated protein kinase (MAPK) pathways along with genes in the *Dusp* family, *Map3k8, Il1b*, and *Il1r2*, among others. However, at 1.5 dpc we observed a strong shift to an anti-inflammatory condition in the IU region. These observations were absent in 0.5 and 1.5 dpp, suggesting that the DEGs observed for those inflammatory responses during pregnancy were likely induced by the presence of sperm. scRNA-seq analysis revealed that the inflammatory responsive genes were likely produced by secretory epithelial cells, compared to other cell types in the oviduct. In addition, multiple DEGs involved in pyruvate and glycolysis were enriched in the IU region, which could provide metabolic support for developing embryos. Lastly, we have also identified that there were cells that express immune markers in the oviduct, indicating that the oviduct is an immuno-dynamic tissue. In conclusion, our findings indicate that the oviduct is adaptive and responsive to the presence of sperm and embryos in a spatiotemporal manner. In this report, we intend to disseminate our findings on the transcriptional profiles during different stages of pregnancy. The complete study and validation at the protein level are currently underway and will be updated as soon as the data are available.

## INTRODUCTION

Optimal physiological conditions in the oviduct (fallopian tube in humans) provide an adaptive microenvironment for several reproductive processes ranging from sperm capacitation and transport, fertilization, and embryonic transport and development, among others [1-10]. With the rise in assisted reproductive technologies (ARTs) however, the oviduct’s influence on gametes has become increasingly underappreciated since embryos can be produced utilizing ARTs *in vitro*. A systematic review of worldwide trends in ARTs from 2003-2014 and a clinical review of ART literature exhibit that nations performing high volumes of ART cycles cannot agree on what methodologies to employ at the clinical level [11,12]. Studies to identify what specific manipulations in ARTs are responsible for producing adverse effects are ongoing [13]. Yet, no definitive conclusions have been elucidated, leaving gaps in our understanding mechanistically to standardize methodologies to produce viable embryos both *in vivo* and *in vitro*.

The oviduct is comprised of four main regions: infundibulum (responsible for oocyte pick-up), ampulla (site of fertilization), isthmus (sperm capacitation/transport and preimplantation embryonic development), and the uterotubal junction (UTJ; where the embryos transit to the uterus). Several studies demonstrated that distal (infundibulum and ampulla: IA) and proximal (isthmus and UTJ: IU) regions of the oviduct have distinct transcriptional profiles [14-17]. However, it is unclear how the presence of the gametes and embryos alter the oviductal responses. The presence of gametes and embryos has been shown to alter gene expression in secretory and ciliated cells of the oviduct during the preimplantation period [18-21]. Additionally, it has been reported that the endometrium responds differently to *in vivo* derived embryos compared to embryos derived utilizing IVF or somatic cell nuclear transfer in large animal models [22,23]. Lastly, there is a growing consensus based on evidence from several species that epigenetic events in preimplantation embryos contribute to altered developmental potential both early and later in life [13]. Indeed, variation in the relative abundance of sets of genes involved in compaction and cavitation, desmosomal glycoproteins, metabolism, mRNA processing, stress, trophoblastic function, and growth and development have been observed in *in vitro* produced embryos when compared to their *in vivo* counterparts [24-27]. However, little is known about the suggested crosstalk and what effect the embryo has on the oviductal environment to modulate early embryo development in mammalian species.

Reciprocal embryo-oviduct interactions stem largely from investigating oviductal transport of fertilized/unfertilized embryos/oocytes in livestock [18,19,28-41] and rodents [14,20,42,43]. It has been suggested that fertilized embryos produce prostaglandin E2 that facilitates transport to the uterus in mares [34,36], whereas non-fertilized embryos remain in the oviduct [28]. In hamsters, fertilized embryos are transported more expeditiously to the uterus compared to unfertilized eggs [42]. In rats, transferred advanced stage embryos (4-cell vs 1-cell) arrive in the uterus prematurely [43]. In pigs [19] and cows [18], pro-inflammatory responses in the oviduct are down regulated by the presence of embryos, suggesting that the embryo may facilitate maternal-embryo tolerance during its passage through the oviduct. Moreover, in humans, embryo-derived platelet-activating factor has been implicated in the control of embryo transport to the uterus [44]. However, in mono-ovulatory species, alterations in the oviductal transcriptome are difficult to detect [33,36] suggesting that the effect of the embryo is localized. Therefore, to amplify transcriptome alterations in heifers, transfer of multiple (up to 50) embryos into the oviduct is necessary [18].

Based on this premise, our study aims to elucidate the adaptive or passive nature of the oviduct during natural fertilization and preimplantation embryo development. We dissected oviducts from naturally fertilized/pseudopregnant mice at 0.5, 1.5, 2.5, and 3.5-days post coitus or pseudopregnancy (dpc or dpp, respectively). Gene expression profiles were analyzed from two different regions of the oviduct: infundibulum + ampulla (IA) and isthmus + UTJ (IU) using bulk-RNA and single cell RNA (scRNA) sequencing analyses. Our RNA expression profile from bulk RNA-seq findings were confirmed by scRNA-seq analysis. In addition, cell-type specific expression of genes of interest were also identified using scRNA-seq analysis. Overall, we observe a critical transition of transcripts in the oviduct from 0.5 dpc to other timepoints that induces large sets of differentially expressed genes (DEGs) in both IA and IU regions. One of our key observations is that an elevated pro-inflammatory transcriptional profile at 0.5 dpc was likely due to the presence of sperm and that a strong anti-inflammatory condition correlated with the presence of the embryo in the IU region.

## MATERIALS AND METHODS

### Mouse handling

All animals were maintained at Washington State University and were handled according to Animal Care and Use Committee guidelines using approved protocols 6147 and 6151. C57BL/6J mice from Jackson Laboratories were used in all experiments in this study. C57BL/6J female mice between 8-16 weeks were naturally mated with fertile C57BL/6J males. Pseudopregnancy was induced by mating females with vasectomized males. The presence of copulatory plug the next morning was considered 0.5 days post coitus (dpc) for females mated with fertile male and 0.5 days of pseudopregnancy (dpp) for females mated with vasectomized males.

### Tissue collection

Oviductal tissues were collected and stored in pairs (one pair of oviducts per animal) at 0.5, 1.5, 2.5, and 3.5 dpc/dpp of natural fertilization and pseudopregnancy. For 0.5 dpc/dpp tissue, female mice were placed for mating at 9 p.m. For 1.5, 2.5, and 3.5 dpc/dpp, female mice were placed for mating between 5-6 p.m. Oviducts were dissected and kept in 1 mL Leibovitz-15 (L15, Gibco, 41300070, ThermoFisher Scientific, Carlsbad, CA) + 1% fetal bovine serum (FBS, Avantor 97068-091, Radnor Township, PA) media. Before sectioning the oviduct into two regions (Infundibulum + Ampulla (IA), Isthmus + Uterotubal Junction (IU), oviducts were flushed with L15 + 1% FBS media under a 37°C dissecting microscope (Leica MZ10f, Leica Microsystems, Buffalo Grove, IL). The presence of a minimum of 6 embryos per female was confirmed as a benchmark to represent the average litter size. Additionally, embryos were confirmed to be in the correct developmental stage and location in oviductal tissue samples. Then, oviducts were sectioned into IA and IU regions. Tissue samples were placed in a sterile Eppendorf tube and flash frozen in liquid nitrogen. Samples were stored at -80°C for later RNA extraction. All dissections took place between 10:00-13:00 h to decrease sample variation. The average time from cervical dislocation of the mouse to flash freezing tissues was 15:43 (min:sec). For dpp samples, the oviducts were collected at the same time points as their dpc counterparts.

### Bulk RNA isolation, sequencing, and analysis

Both tissue and embryo total RNA were extracted utilizing the RNeasy Micro Kit (Qiagen, Germantown, MD) according to the manufacturer’s instructions. A DNA digestion was performed with all samples using Qiagen RNase-free DNase Set (1500 K units). RNA was then shipped to University of California San Diego (UCSD) for quality control, library preparation, and sequencing. RNA integrity (RIN) was verified using TapeStation for a minimum RIN value of 7. RNA from this study has an average RIN of 9.04. RNA libraries were prepared using Illumina Stranded mRNA Prep kit (Illumina Inc., San Diego, CA). Then libraries were sequenced using the Illumina NovaSeqS4 platform with a read depth of 25M reads/sample (*n*=3/region/timepoint), paired-end, and 100bp read length. FASTQ files were then analyzed utilizing BioJupies [45] and integrated web application for differential gene expression and pathway analysis (iDEP) [46]. The quality control, sequence alignment, quantification, differential gene expression (DEG), heatmaps, and pathway and enrichment analyses were performed using default settings as indicated in BioJupies and iDEP webtools [45,46]. In brief, FASTQ files were pseudoaligned and DEGs were determined using DESeq. DEGs were then plotted as Principal Component Analysis (PCA) and heat maps through BioJupies. In some cases, read counts or reads per kilobase of transcript per million mapped reads (FPKM) were exported from BioJupies and imported into iDEP.92 for further pathway and KEGG analyses. Venny 2.1 [47] was used to generate common/overlap gene lists between different regions and timepoints.

To validate that our isolation method and RNA-seq data analysis pipeline are reproducible with the previous report [14] we evaluated the gene expression profiles of IA and IU regions from estrus samples (n=3 mice/region). In agreement with the previous findings [14], principal component analysis (PCA) plots showed that the IA and IU regions segregate from each other along the PC1 axis (74.3%) with respect to estrus (data not shown). Similar to the previous report, there was a significant indication of a region-specific expression of large subsets of genes.

### Single cell isolation, library preparation and single cell RNA-sequencing (scRNA-seq)

Another set of mice were used for single cell isolations and scRNA-seq analysis. Mating and tissue collection protocols were similar to bulk RNA isolation described above with an exception that female mice were superovulated using protocol described previously [48] to ensure sufficient numbers of female mice at each time point could be harvested for single cell isolation and library preparation within the same day (n= 5-6 mice/group). Superovulation was performed by intraperitoneal injection of 5 IU pregnant mare serum gonadotropin (PMSG, Prospect HOR-272, East Brunswick, NJ). Forty-eight hrs after PMSG injection, females were injected with 5 IU of human chorionic gonadotropin (hCG, Prospect HOR-250). Immediately after hCG injection, females were placed in fertile male cages for mating. Oviducts were collected at 0.5, 1.5, and 2.5 dpc and dissected into IA and IU regions before single cell isolation. For the control group, oviducts were collected 16 hrs post hCG injection. Trypsin-EDTA (0.25%, MilliporeSigma, T4049) was used for oviductal cell isolation using our previously described method [15]. The final cell concentration was targeted for 8,000 cells/run. Cell singlets were captured for the library preparation using 10X Chromium Controller and Chromium Next GEM Single Cell 3’ GEM, Library & Gel Bead Kit v2 (10X Genomics, Pleasanton, CA). Libraries generated were then evaluated for quality using Fragment Analyzer (Agilent, Santa Clara, CA). Libraries were sequenced using Illumina HiSeq4000 at University of Oregon, targeting 400 million reads/run, paired-end, and 100 bp read length. scRNA-seq output for each dataset is listed in Table 1.

**Table 1:** scRNA-seq output for each dataset in this study

| Sample Name | Estimate number of cells | Total reads | Mean reads/cell | Median gene/cell | Median UMI counts/cell | Total detected gene # | Sequencing saturation |
| --- | --- | --- | --- | --- | --- | --- | --- |
| Ctrl_IA | 8,793 | 89M | 10,153 | 1,387 | 2,976 | 19,843 | 32.1% |
| Ctrl_IU | 10,387 | 95M | 9,173 | 967 | 2,194 | 19,285 | 36.6% |
| 0.5_IA | 3,689 | 105M | 28,382 | 1,585 | 3,891 | 19,044 | 64.4% |
| 0.5_IU | 7,551 | 93M | 12,312 | 731 | 1,562 | 18,612 | 67.1% |
| 1.5_IA | 11,760 | 84M | 7,184 | 1,138 | 2,116 | 19,941 | 28.0% |
| 1.5_IU | 10,480 | 84M | 7,989 | 860 | 1,792 | 19,525 | 35.6% |
| 2.5_IA | 4,504 | 107M | 23,870 | 1,616 | 3,590 | 19,250 | 60.5% |
| 2.5_IU | 7,804 | 100M | 12,871 | 928 | 2,178 | 18,350 | 65.0% |

### scRNA-seq analysis

Scanpy was used to analyze the scRNA-seq data. The generated loom files were read in as anndata objects and concatenated into a master anndata object. Preprocessing and quality control were performed similarly to the methods described in Scanpys clustering tutorial [49] and Seurat’s clustering tutorial [50]. Filtered out were cells expressing fewer than 200 genes, genes expressed in fewer than 3 cells, doublets (cells/droplets with counts for greater than 4,000 genes), and cells with greater than 5% mitochondrial gene counts. Total counts were normalized to 10,000 for every cell and log transformed. Highly variable genes were then identified using scanpy’s “highly_variable_genes” function with default parameters. Effects of mitochondrial gene expression and total counts were regressed out, and the data scaled to unit variance and a mean of zero. Dimensionality reduction was first achieved through principal component analysis with Scanpy’s default parameters. To achieve further dimensionality reduction, a neighborhood graph of cells was computed, utilizing the top 40 principal components (PCs) and a neighborhood size of 10, then embedded utilizing Uniform Manifold Approximation and Projection (UMAP), using the default parameters in Scanpy. Clustering of cells was achieved through Leiden clustering at a resolution of 0.1. Established marker genes were used to identify clusters as specific cell types: pan-epithelial (*Epcam*+), secretory (*Pax8*+), ciliated (*Foxj1*+), leukocytes (*Ptprc*+), antigen-presenting cells (*Cd74+*), monocytes and macrophages (*Ms4a7+ Cd14+*), T-cells (*Cd3d+, Cd3g*), natural killer and NKT cells (*Nkg7+, Klrb1c+*), B-cells (*Cd79a+, Cd79b+*), granulocytes (*S100a8+, S100a9+*), Neutrophils (*Ly6g+*), fibroblasts and stromal (*Pdgfra*+, *Twist2*+, *Dcn*+, *Col1a1*+), and endothelial (*Pecam1*+). Subsets containing only specific cell types (e.g., secretory cells), treatments (e.g., control, 0.5, 1.5, and 2.5 dpc), or regions (e.g., IA and IU) were created for specific downstream analyses and analyzed through the same process as above with identical parameters. Source code for these analyses and figure generation will be available at GitHub upon publication of this manuscript.

### Gene ontology (GO)

Differentially expressed genes were identified using scanpy’s “highly_variable_genes” function with default parameters. Differentially expressed gene lists were generated during bulk and single-cell RNA-seq data analysis as described in their respective methods sections. Generated DEG sub lists containing up- and downregulated genes with a log2FC ≥ 1 or ≤ -1 respectively were then filtered for genes. The filtered gene lists were submitted to Enrichr for Gene Ontology enrichment analysis. Exported data were plotted in R using ggplot [51].

### Data availability

Raw data as fastq files were deposited at Gene Expression Omnibus and will be publicly available upon publication of this manuscript.

### Histological analysis

Another set of mice were used for histological analysis. Procedures for mating and oviduct collection were similar to those for bulk-RNA-seq procedures. Tissue samples for 0.5, 1.5, 2.5, and 3.5 dpc were formalin-fixed, paraffin-embedded and were sectioned at a 5 µm thickness. Some sections were stained with hematoxylin and eosin (H&E) using standard staining procedure as previously described [15].

## RESULTS

### Oviductal transcriptomics are dynamic during pregnancy and pseudopregnancy

First, to confirm the presence and location of embryos in the oviduct in our mouse model, we sampled the oviduct at different timepoints and evaluated the location of the embryos using H&E staining. Fertilized embryos were not present at the infundibulum but located in the ampulla at 0.5 dpc (**Fig. 1**). Two-cell embryos were located in the isthmus at 1.5 dpc. Embryos were then halted at 2.5 dpc in the uterotubal junction at the 8-cell stage up until the morula stage. At 3.5 dpc, the UTJ region was devoid of embryos as all embryos were transported to the uterus.

**Fig. 1.**
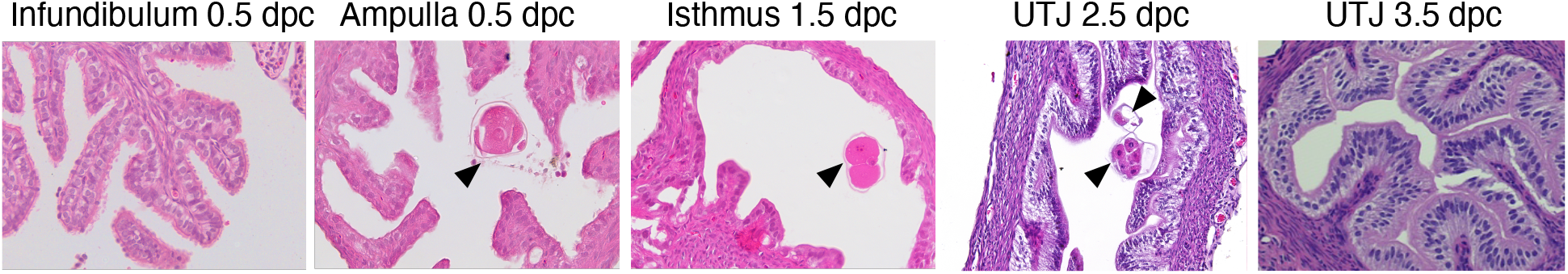
Histological analysis of different oviductal regions (infundibulum, ampulla, isthmus, and uterotubal junction (UTJ) in mice at different stages during preimplantation embryo development (0.5, 1.5, 2.5, and 3.5-days post coitus or dpc) using H&E analysis. Arrowheads = embryos. Representative images are shown at 200x magnification (n=3 mice/timepoint/region).

Next, to determine whether the transcriptional profiles of each oviductal region are unique at fertilization and different developmental stages in preimplantation development, bulk RNA-seq analysis was performed at 0.5, 1.5, 2.5, and 3.5 dpc. Additionally, we aim to address whether changes in transcriptional signatures in the oviduct are governed by hormonal fluctuations or the presence of sperm/embryos. Therefore, pseudopregnant oviducts at corresponding stages (0.5, 1.5, 2.5, and 3.5 dpp) were used for comparisons. PCA plots were generated with respect to the top 2,500 DEGs (**Fig. 2**). Surprisingly, with respect to both the IA and IU regions, overall transcripts at 0.5 dpc (**Fig. 2A**) and 0.5 dpp (**Fig. 2B**) were segregated to the right most axis, while 1.5-3.5 dpc and dpp biological replicates were either segregated to the left axis or transitioning to the left axis. These data suggest that transcripts at 0.5 dpc and dpp in the oviduct are unique and differ from those at 1.5-3.5 dpc and dpp.

**Fig. 2.**
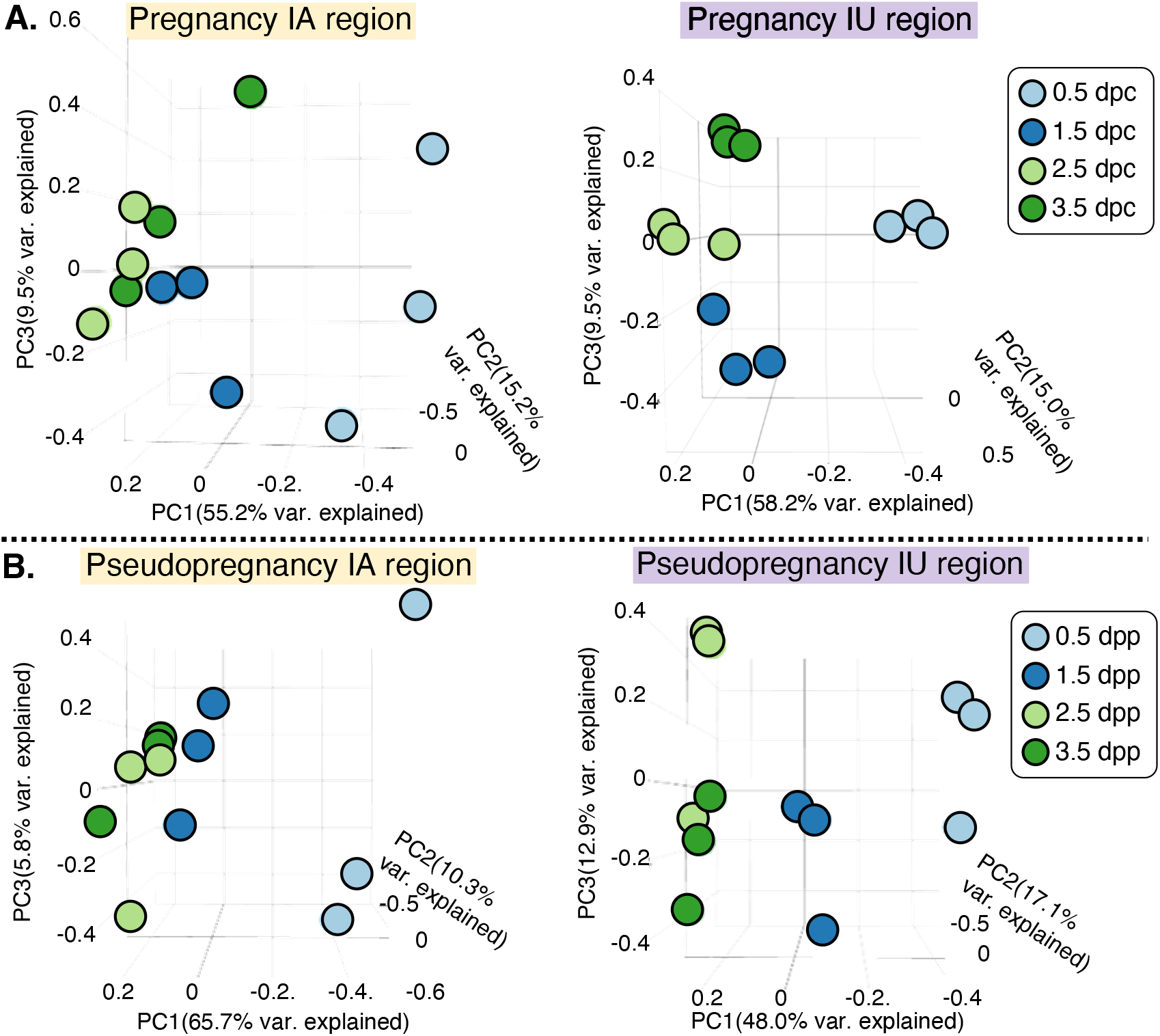
Principal Component Analysis of top 2500 DEGs identified from bulk-RNA seq of the oviduct collected during **A**. pregnancy at 0.5, 1.5, 2.5, and 3.5 days of pregnancy (dpc) and **B**. pseudopregnancy at 0.5, 1.5, 2.5, and 3.5 days of pseudopregnancy (dpp). Data are separated into infundibulum-ampulla (IA) compared to isthmus-uterotubal junction (IU) regions (n=3 mice/timepoint/region).

Before we specifically determine unique genes in the oviduct at each timepoint during pregnancy, it is noted that expression signatures of the top 2500 DEGs in the IA region during pseudopregnancy were similar to those during pregnancy as indicated by a heatmap generated using unsupervised hierarchical clustering (**Fig. 3A)**. However, there were exceptions at 0.5 (**Fig. 3A**, blue box) and 1.5 (**Fig. 3A**, black box) dpc vs. dpp. Unlike the IA region, DEGs in the IU region were more dynamic as indicated by the presence of unique sets of genes at 0.5 (**Fig. 3B**, blue box), 1.5 (**Fig. 3B**, black box), 2.5 (**Fig. 3B**, red boxes), and 3.5 (**Fig. 3B**, purple box) timepoints between pregnancy vs pseudopregnancy. Overall, data suggest that the transcriptional profile in the oviduct at all stages during the preimplantation period in the IU region is more dynamic compared to the IA region.

**Fig. 3.**
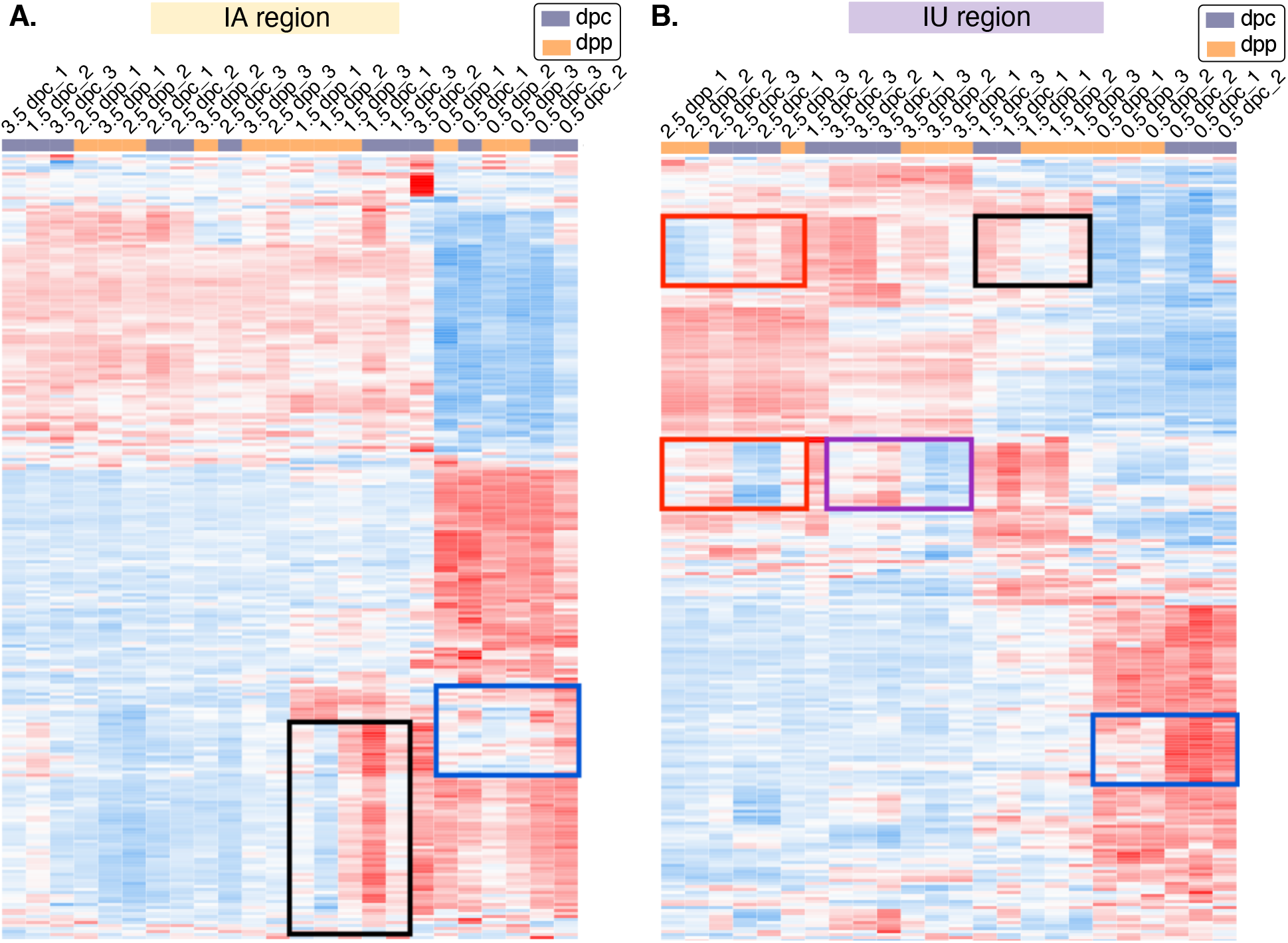
Heatmap plots of unsupervised hierarchical clustering of top 2500 DEGs in the oviduct during pregnancy (0.5, 1.5, 2.5, and 3.5 dpc) compared to pseudopregnancy (0.5, 1.5, 2.5, and 3.5 dpp) of **A**. IA and **B**. IU regions. Gene expression patterns that differ between 0.5 dpc (blue boxes), 1.5 dpc (black boxes), 2.5 dpc (red boxes), and 3.5 dpc (purple box) compared to the corresponding dpp (n=3 mice/timepoint/region). Blue; downregulated, red; upregulated.

As we observed that oviductal transcription signatures were more unique at 0.5 when compared to those at 1.5-3.5 dpc (i.e., 0.5 dpc vs. rest) in both IA (**Fig. 4A**) and IU (**Fig. 4B**) regions, we next determined biological processes of genes that were differentially expressed at 0.5 dpc compared to 1.5-3.5 dpc in both IA and IU regions. This analysis (0.5 dpc vs rest) was chosen to isolate what general processes may be occurring during the transition from 0.5 dpc to the other time points. We found that there were no significant GO biological processes evaluated from the downregulated DEGs lists (adjusted *p*-value > 0.05) at 0.5 dpc compared to 1.5-3.5 dpc in the IA or IU regions. However, unique DEGs in the IA region that were upregulated at 0.5 dpc compared to other timepoints were enriched for the following biological processes: extracellular matrix (ECM) organization, extracellular structure organization, collagen fibril organization, and Ca2+ ion homeostasis, among others (**Fig. 4C**). Therefore, it is possible that transcripts in IA region adapted to the presence of cumulus-oocyte complexes (COCs) and required expansion of the mucosal folds to accommodate a large volume of COCs, especially in the ampulla region (**Fig. 1**). Most interestingly however, we found upregulated DEGs enriched for biological processes at 0.5 dpc in the IU region including cellular response to cytokine stimulus, neutrophil migration, response to interferon-gamma, response to lipopolysaccharide, and neutrophil chemotaxis, (**Fig. 4D**). Moreover, there are multiple biological processes involved in glucose catabolic process to pyruvate and pyruvate metabolic processes that were uniquely upregulated at 0.5 dpc (and subsequently downregulated during 1.5-3.5 dpc) in the IU region (**Fig. 4D**). Therefore, it is likely that, at 0.5 dpc, the isthmus region of the oviduct was heavily regulated for an inflammatory response in the presence of sperm while simultaneously preparing for the metabolic switch of the embryos from pyruvate to glucose metabolism.

**Fig. 4.**
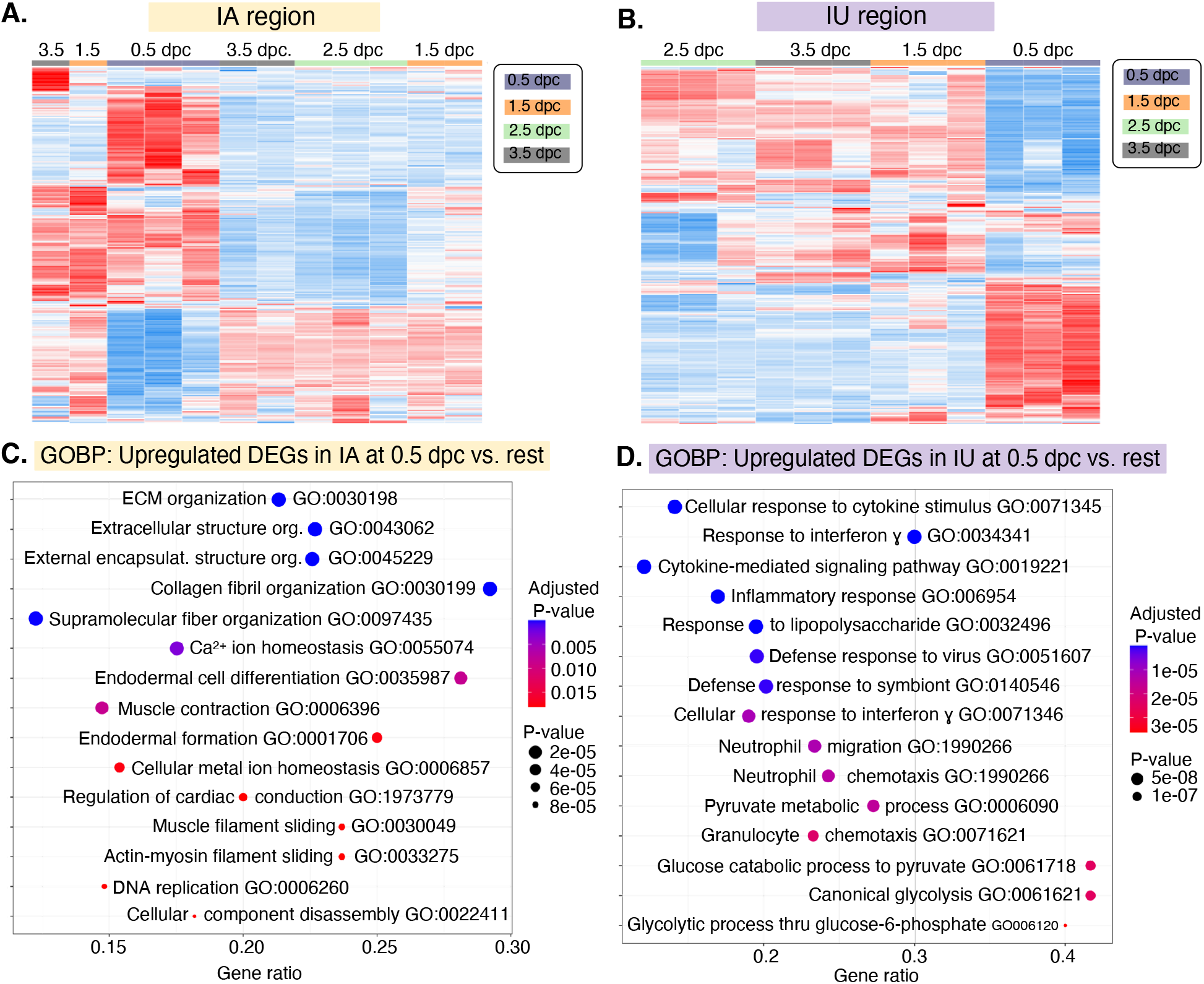
Heatmap plots of unsupervised hierarchical clustering of top 2500 DEGs identified from bulk-RNA seq in the oviduct during pregnancy (0.5, 1.5, 2.5, and 3.5 dpc) of **A**. IA and **B**. IU regions. Dot Plots of biological processes from Gene Ontology (GOBP) analysis from DEGs that were upregulated at 0.5 dpc compared to 1.5-3.5 dpc (0.5 dpc vs. rest) in **C**. IA and **D**. IU regions. GO numbers were indicated within graphs. ECM; extracellular matrix, Org; organization. Gene ratio indicates the ratio of genes that were present in our dataset compared to all genes in the database for each BP.

### Oviductal transcriptomes that are unique to the distal (IA) region in the presence/absence of sperm

It is also important to note that COCs are present in the IA region in both the pregnant and pseudopregnant samples. Therefore, the only difference would be the presence of sperm in the IA region at 0.5 dpc, while the IA region at 0.5 dpp lacks sperm. We found that upregulated DEGs at 0.5 dpc compared to 0.5 dpp demonstrated a significant increase in biological processes that were involved in cytokine-mediated signaling pathway, inflammatory response, regulation of phagocytosis, and cellular response to type I interferon, among others (**Fig. 5A**). This finding recapitulates the data from 0.5 dpc at IU region (**Fig. 4D**), at which was exposed to sperm. Therefore, these results reinforce the hypothesis that sperm is the major stimulator for the observed inflammatory response in the oviduct in both IA and IU regions.

**Fig. 5.**
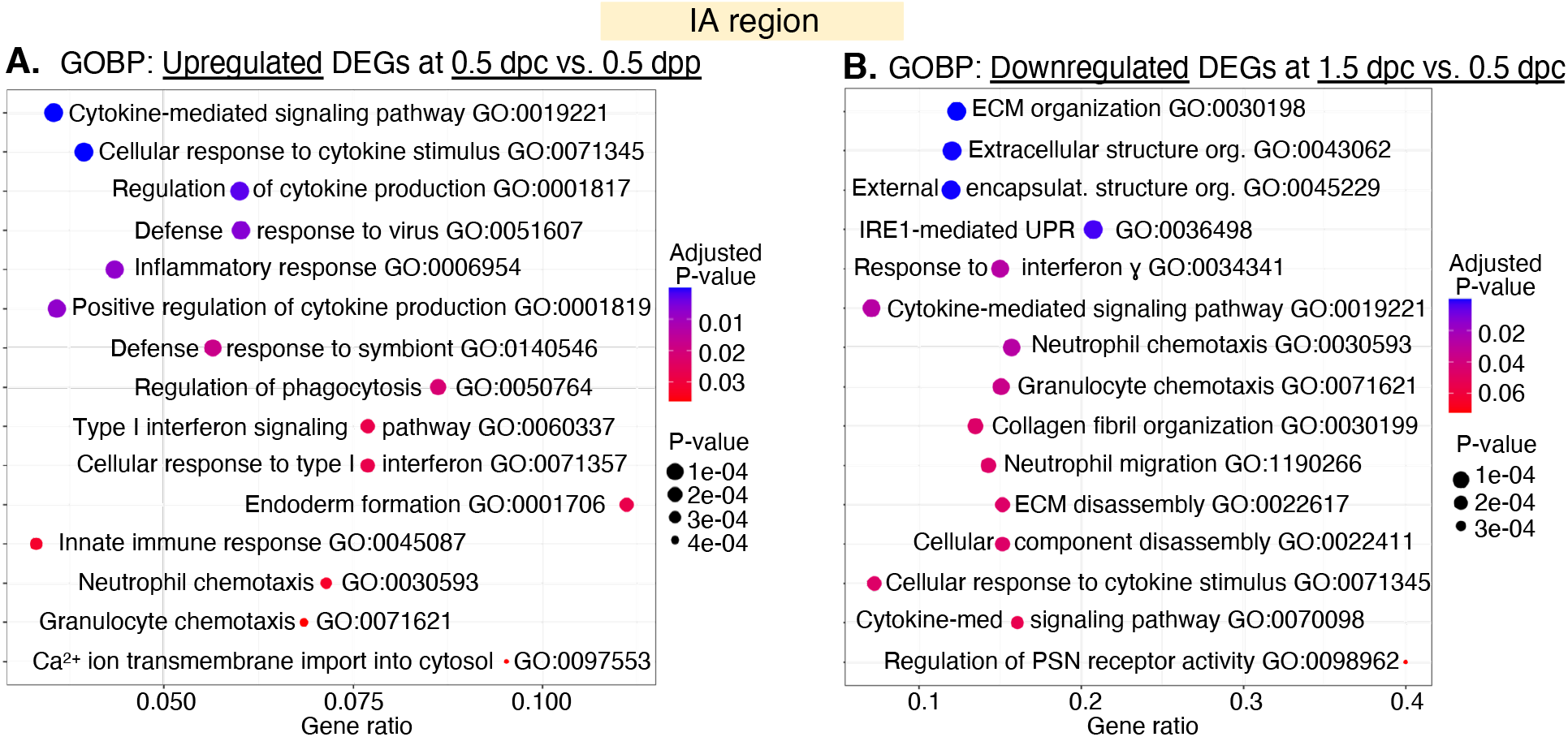
Dot Plots of biological processes from Gene Ontology (GOBP) analysis from DEGs identified from bulk-RNA seq were **A**. upregulated at 0.5 dpc compared to 0.5 dpp in the IA region and **B**. downregulated at 1.5 dpc compared to 0.5 dpc in the IA region. GO numbers were indicated within graphs. ECM; extracellular matrix, Org; organization, IRE1; Inositol-requiring enzyme 1, UPR; unfolded protein response; med; mediated, PNS; postsynaptic neurotransmitter. Gene ratio indicates the ratio of genes that were present in our dataset compared to all genes in the database for each BP.

Next, we evaluated the IA region at 0.5 dpc compared to 1.5 dpc (presence of sperm vs. post-sperm exposure, respectively). There were no significant (adjusted *p*-value > 0.05) GO pathways from upregulated DEGs in the IA region at 1.5 dpc. However, we observed significant biological processes that were enriched for downregulated DEGs at the IA region at 1.5 dpc compared to 0.5 dpc. These processes included response to interferon-gamma, neutrophil chemotaxis, cytokine-mediated signaling pathway, and neutrophil migration (**Fig. 5B**). In addition to downregulated DEGs involved in inflammatory pathways, there were several biological processes identified at 1.5 dpc (**Fig. 5B**). These additional biological processes were common (but DEGs were expressed in the opposite direction, i.e., downregulated at 1.5 dpc vs. upregulated at 0.5 dpc) to the IA region at 0.5 dpc (**Fig. 4C**), such as ECM organization, extracellular structure organization, and collagen fibril organization. Therefore, it is possible that transcripts in the IA region at 1.5 dpc appear to revert back to the environment before COCs and sperm exposure. These findings suggest that the IA region is dynamic, albeit less so when compared to the IU region.

### Oviductal transcriptomes that are unique to the proximal (IU) region in the presence or absence of sperm and embryos

As highlighted above, DEGs were more dynamic in the IU region at different timepoints (**Fig. 4B**) during preimplantation embryo development compared to that of the IA region. At 0.5 dpc, the sperm are present, creating a sperm reservoir in the isthmus [52]. There were no significantly downregulated processes (adjusted *p*-value > 0.05) at 0.5 dpc when compared to 0.5 dpp. However, when comparing 0.5 dpc to 0.5 dpp in the IU region (presence or absence of sperm, respectively), GO analysis revealed enrichment of multiple proinflammatory biological processes, including inflammatory response, neutrophil migration, neutrophil chemotaxis, positive regulation of acute inflammatory response, and response to lipopolysaccharide when sperm were present in the IU (**Fig. 6A**). Subsequently, we performed a similar analysis comparing 0.5 dpc to 1.5 dpc of pregnancy in the IU region (presence of sperm vs. post-sperm exposure). There were no significantly upregulated processes at 1.5 dpc when compared to 0.5 dpc. Here, the most striking results were observed when we evaluated the biological processes for downregulated DEGs at 1.5 dpc when compared to 0.5 dpc. We found that the majority of these biological processes were cellular response to cytokine stimulus, response to interferon gamma, response to lipopolysaccharide, and neutrophil chemotaxis (**Fig. 6B**). These data suggest that the oviduct may suppress the response to inflammation in the isthmus once the sperm are clear in order to become conducive for the embryo’s survival at 1.5 dpc.

**Fig. 6.**
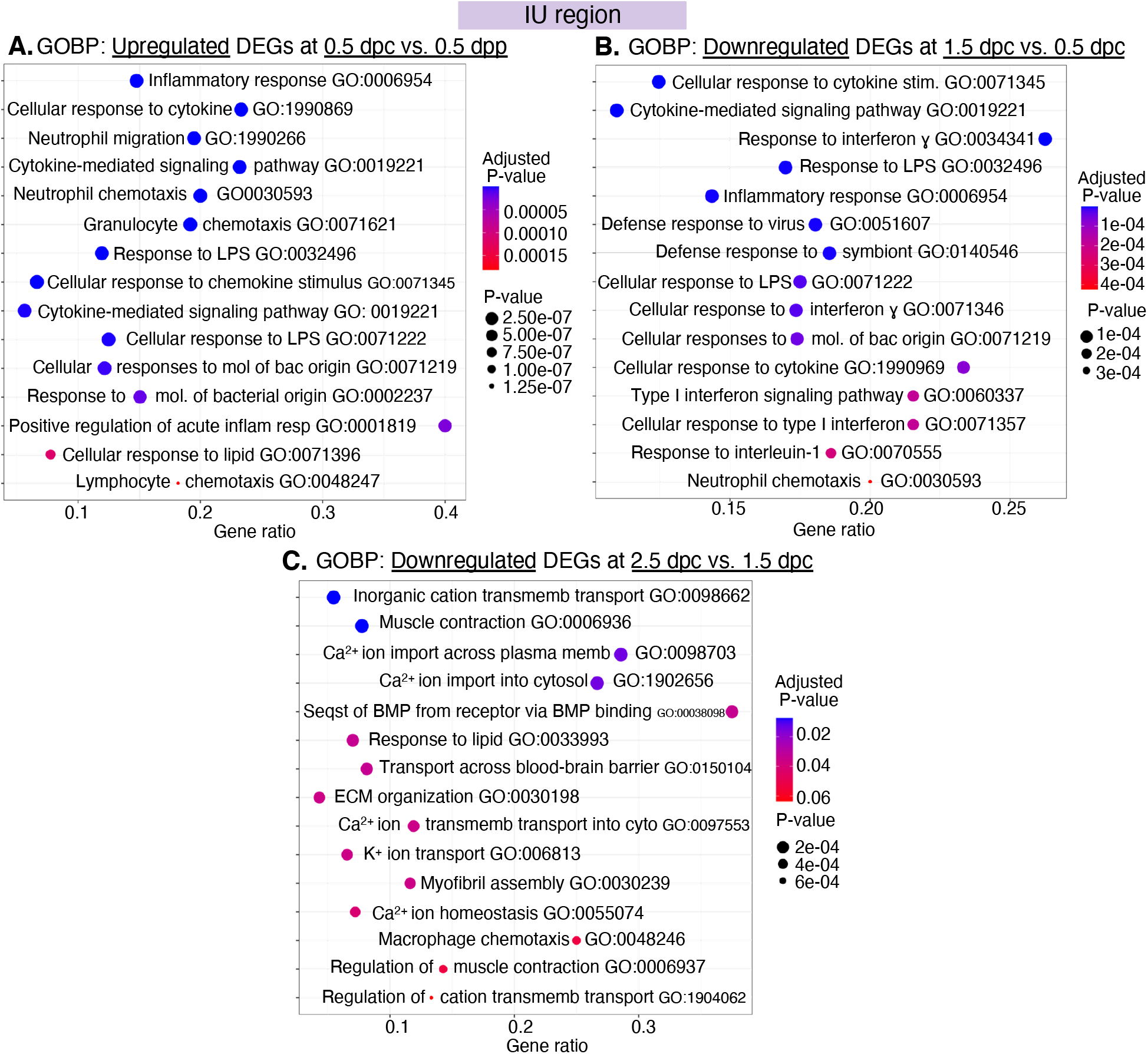
GOBPs dot plots of DEGs identified from bulk-RNA seq in the IU regions when compared between **A**. upregulated DEGs at 0.5 dpc compared 0.5 dpp, **B**. downregulated DEGs at 1.5 dpc compared to 0.5 dpc, and **C**. downregulated DEGs at 2.5 dpc compared to 1.5 dpc. GO numbers were indicated within graphs. LPS; lipopolysaccharide, mol; molecular, bac; bacterial; inflam resp; inflammatory response, transmemb; transmembrane, memb; membrane, seqst; sequestering, BMP; bone metalloprotease, cyto; cytosol. Gene ratio indicates the ratio of genes that were present in our dataset compared to all genes in the database for each BP.

At 1.5-2.5 dpc, 2-cell through morula/blastocyst stage embryos move back and forth within the isthmus [53]. We found that there were no significant biological processes enriched from upregulated DEGs at 2.5 dpc when compared to 1.5 dpc at the IU region. However, GO analysis of downregulated DEGs at 2.5 dpc compared to 1.5 dpc reveals the downregulation of genes involved in sequestering of BMP from receptor via BMP binding, regulation of muscle contraction, Ca^2+^ ion import across plasma membrane, and inorganic cation transmembrane transport (**Fig. 6C**). Lastly, there were no significant biological processes identified from neither upregulated or downregulated DEGs at 3.5 dpc when compared to 2.5 dpc.

### scRNA-seq reveals that secretory epithelial cells contribute to the pro- and anti-inflammatory responses in the oviduct

To evaluate the cell-type specific transcripts, scRNA-seq analyses were used. Embryos are present in the oviduct during preimplantation periods at 0.5, 1.5, and 2.5 dpc. Since we did not observe significant transcriptional changes from bulk RNA-seq dataset at 3.5 dpc, we opted not to assess a 3.5 dpc data point in our scRNA-seq analysis. In this experiment, superovulated, non-mated samples were used as a control (Ctrl). First, we confirmed that all cell types are present in the oviduct (**Fig. 7A**) similar to what was previously reported [15], with some exceptions for immune cell types (see below). There was minimum overlap between cells isolated from IA or IU regions (**Fig. 7B**). In addition, all cell types were present at all time points except for an *Ephx2*+ cluster (only present at 0.5 dpc and Ctrl) and a neutrophil cluster (*Ly6g+*, only present at 0.5 dpc) (**Fig. 7C**). We then investigated whether our findings from our bulk RNA-seq data would be recapitulated in the scRNA-Seq dataset. Here, we exclusively focused on the IU region as it is the most dynamically regulated during pregnancy. We observed a similar pro- and anti-inflammatory pattern in our scRNA-seq pregnancy replicates compared to the bulk RNA-seq data (**Fig. 7D**).

**Fig. 7.**
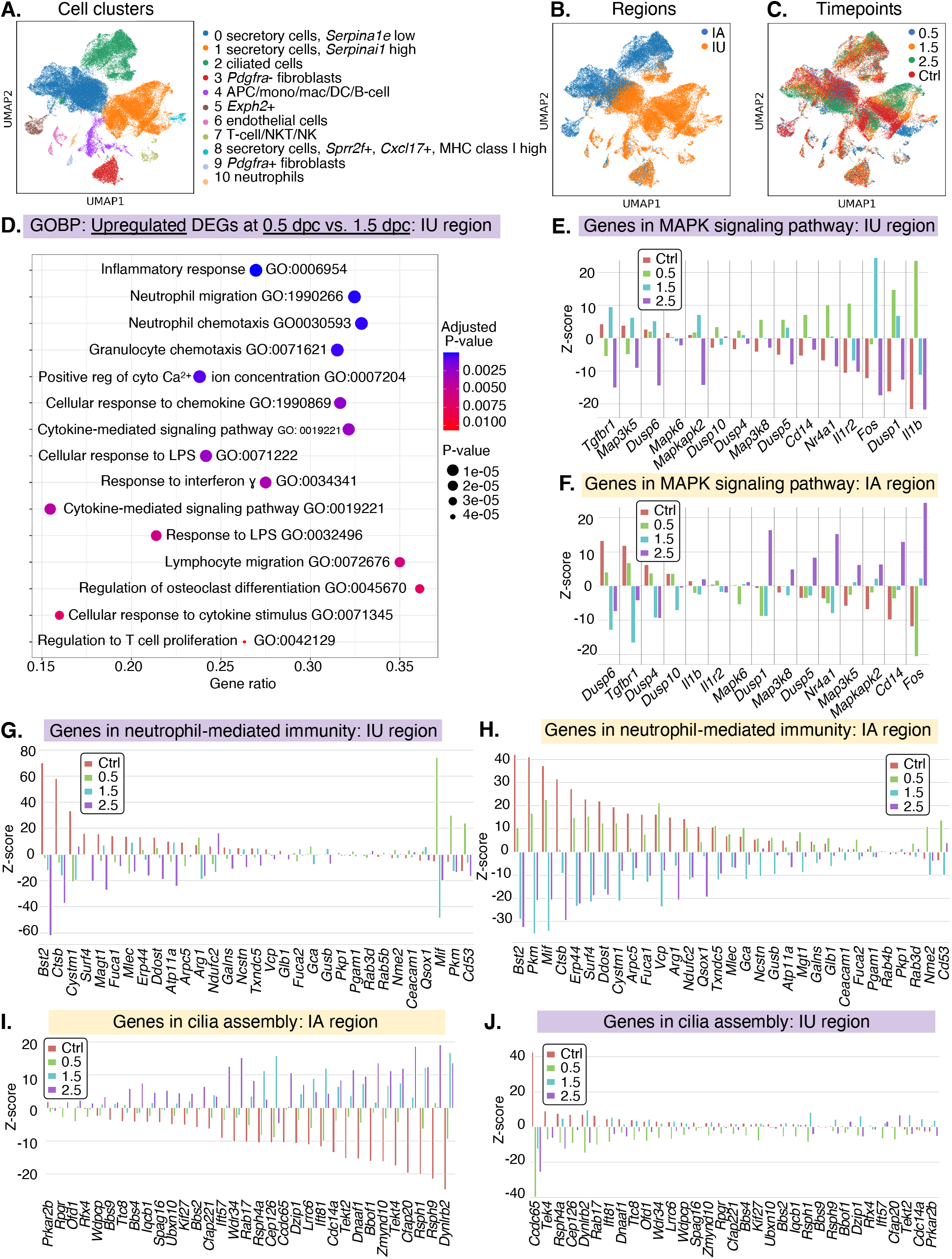
scRNA-seq analysis of the oviduct at 0.5, 1.5, 2.5 dpc and control (Ctrl). Uniform Manifold Approximation and Projection (UMAP) of **A**. cell clusters identified from the oviduct **B**. at different regions (IA and IU) and **C**. at different timepoints (0.5, 1.5, 2.5, and Ctrl). **D**. Dot plots of GOBPs when compared between upregulated DEGs at 0.5 dpc compared to 1.5 dpc from all cell populations in the IU regions. GO numbers were indicated within graphs. LPS; lipopolysaccharide. Gene ratio indicates the ratio of genes that were present in our dataset compared to all genes in the database for each BP. **E.-H**. Z-score bar graphs of genes in (**E**. and **F**.) MAPK signaling pathway as well as (**G**. and **H**.) in neutrophil-mediated immunity, activation, and degranulation at all timepoints in all cell populations at the IU and IA regions. **I**. and **J**. Z-score bar graphs of genes in the cilia assembly in all cell populations at the (**I**.) IA and (**J**.) IU regions at all timepoints.

Based on GO pathway analysis of our bulk RNA-seq finding, we further investigated several genes that were upregulated at 0.5 and 1.5 dpc corresponding to GO terms ‘inactivation of mitogen activated protein kinase (MAPK) activity’ and ‘MAP kinase phosphatase activity’. Genes associated with these pathways were mostly upregulated at 0.5 and 1.5 dpc (**Fig. 7E**, green and teal bars) and downregulated in Ctrl and 2.5 dpc in the IU regions (**Fig. 7E**, red and purple bars). These genes include dual-specificity phosphatase family (*Dusp1, Dusp5, Dusp6, Dusp10*), *Fos, interleukin 1b* (*Il1b*), IL1 receptor 2 *(Il1rb*), and others. Expression patterns of many of these genes were found to be the opposite in the IA region (**Fig. 7F**). We also further assessed several genes from the GO term ‘neutrophil-mediated immunity’ to explain the appearance and subsequent disappearance of the neutrophil cluster at 0.5 and 1.5 dpc, respectively. Interestingly, these genes were found to be downregulated in both the IU and IA regions at the 1.5 and 2.5 dpc timepoints (**Fig. 7G-H**, teal and purple bars).

As it was previously shown that ciliary activity is heavily involved in dispersing the COCs in the ampulla by generating a fluid flow in a circular motion at 0.5 dpc [53], we evaluate the genes involved in ciliated cell function across all timepoints. We found that genes involved in cilia assembly were severely suppressed in Ctrl and at 0.5 dpc in the IA region (**Fig. 7I**, red and green bars) and expression levels of these genes increased at 1.5 and 2.5 dpc (**Fig. 7I**, teal and purple bars). Interestingly, expression of these genes was downregulated at 0.5 and 1.5 dpc (**Fig. 7J**, green and teal bars), while upregulated in the Ctrl and 2.5 dpc (**Fig. 7J**, red and purple bars).

Most importantly, to elucidate which cell population is contributing to both the observed pro- and anti-inflammatory response in both IA and IU regions at 0.5 and 1.5 dpc, we evaluated DEGs in each cell population and performed GO pathway analysis. We found that upregulated DEGs from secretory epithelial cells (both clusters 1 and 2; **Fig 7A**) from both IA and IU regions at 0.5 dpc compared to 1.5 and 2.5 dpc were enriched for the biological processes involved in inflammatory response, neutrophil migration, cellular response to chemokine, chemokine-mediated signaling pathways, and several others chemokine signaling pathways (**Fig. 8A**). In contrast, when evaluated for upregulated DEGs in ciliated epithelial cells, there were no significantly enriched biological processes (**Fig. 8B**, adjusted *p*-value > 0.05). Therefore, these findings from scRNA-seq analysis recapitulate our bulk RNA-seq findings. In addition, it also showed that secretory cells were most likely contributing to the DEGs that were highly enriched for pro- and anti-inflammatory biological pathways compared to ciliated epithelial cells and other cell types identified in our dataset.

**Fig. 8.**
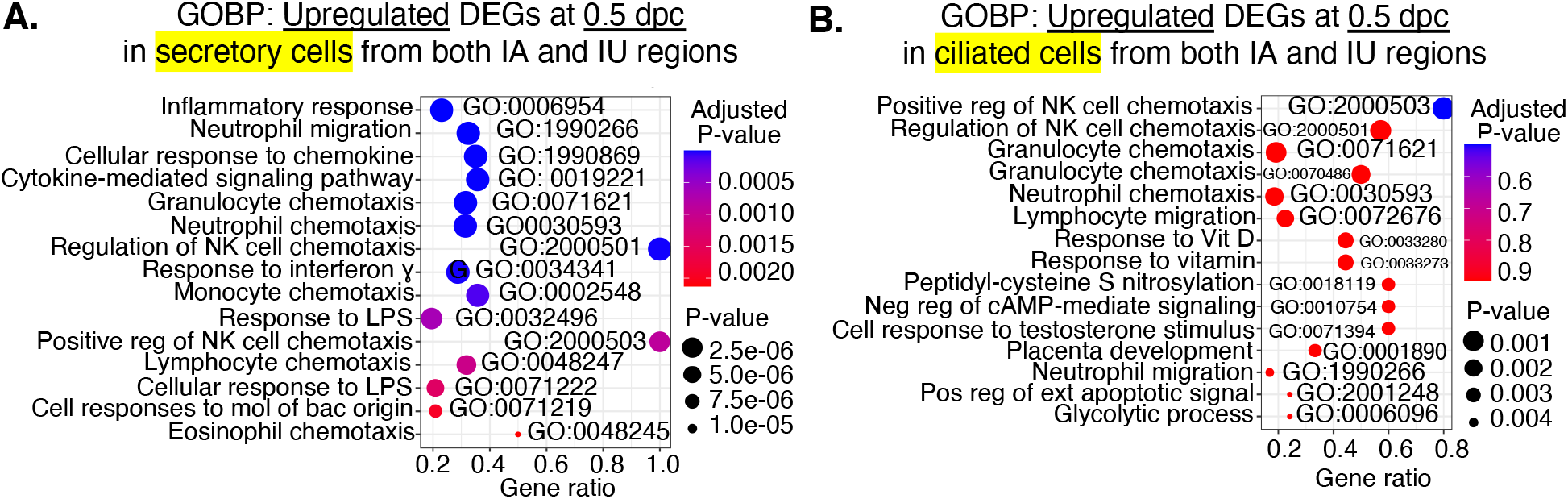
GOBPs dot plots of scRNA-seq analysis when compared between upregulated DEGs from **A**. secretory epithelial cells and **B**. ciliated epithelial cells at 0.5 dpc compared Ctrl from both IA and IU regions. GO numbers were indicated within graphs. NK; natural killer, reg; regulation, neg; negative, cell; cellular, pos; positive, ext; extrinsic. Gene ratio indicates the ratio of genes that were present in our dataset compared to all genes in the database for each BP.

## DISCUSSION

Here we show that the transcriptional profile of the oviduct from both the IA and IU regions are mostly distinct at 0.5 dpc compared to other timepoints based on PCA analyses. This unique transcription pattern was observed at 0.5 in both pregnancy and pseudopregnancy datasets. At this 0.5 timepoint, COCs are present in the IA region in oviducts from pregnant and pseudopregnant females. Large sets of DEGs display a dramatic shift from either being up-or downregulated between 0.5 and 1.5 dpc/dpp. This trend persists for the entirety of the preimplantation development period, with some unique clusters of genes that are dynamic between timepoints 1.5-3.5 dpc. However, the number of dynamic clusters of genes are greater in the IU region than in the IA region. This suggests that the IU region is more dynamic and responsive to the presence of gametes (sperm and oocytes/embryos), whereas the IA region is less dynamic and may be influenced by constituents released from follicular fluid or the physical presence of the COCs, or both.

However, the presence of sperm is mainly unique to the IU region at 0.5 dpc and not during pseudopregnancy. We postulate that sperm are the key mediators in the IU region, stimulating transcriptional changes that are involved in inflammatory responses in the oviduct at 0.5 dpc. At 1.5-3.5 dpc, we found that oviductal transcriptional profiles were more similar to each other compared to that of 0.5 dpc. During this preimplantation developmental period (1.5-3.5 dpc), embryos transit the IA region and arrive in the IU region. It indicates that there could be a critical transition of transcripts from 0.5 dpc/dpp to other timepoints when the embryos are present at 1.5-3.5 dpc. Therefore, our observation suggests that the oviduct is dynamically responsive in a unique manner to the presence of gametes (COCs/sperm) as well as to embryos.

Histological analysis showed that the luminal area in the ampulla region is increased with decreased infoldings to accommodate for the ovulated COCs at 0.5 dpc. At this time point, we found that there were unique upregulated DEGs that corresponded with the biological processes involved in tissue remodeling and muscle filament sliding, such as ECM and collagen fibril organization. Wang and Larina showed that during this timepoint, ciliated epithelial cells in the ampulla region are responsible for creating a circular motion of the COCs within the ampulla [53]. Here, using scRNA-seq analysis we found that the ciliated cell population showed suppressed expression of genes involved in cilia assembly at 0.5 dpc and in Ctrl (COCs are present in these two groups) in the IA region, while almost all genes in cilia assembly were upregulated at 1.5 and 2.5 dpc in the IA region. This finding suggests that when the COCs are present in the ampulla, ciliated cells are functionally active while genes involved in cilia assembly are not being expressed. It is possible that ciliary stalks are physically damaged after such strong motion and physical interaction with COCs at 0.5 dpc, repairing of ciliated cells could be required afterwards, hence the upregulation of genes involved in cilia assembly at 1.5 and 2.5 dpc in ciliated epithelial cells in our scRNA-seq dataset. This finding was reinforced as the pattern of upregulation of cilia assembly at 1.5 and 2.5 dpc was unique to the IA, not the IU region.

At 0.5 dpc, sperm are present in the IU region as a reservoir. Both our bulk RNA-seq and scRNA-seq datasets indicate that sperm are the key influencers for stimulating inflammatory responsive pathways in the oviduct at 0.5 dpc. Specifically, the presence of sperm at 0.5 dpc induces a strong pro-inflammatory response in the IU region with upregulation of genes involved in inflammatory cytokines, neutrophil activation, lymphocyte recruitment and T-cell proliferation. These findings were consistently captured from both bulk RNA-seq and scRNA-seq datasets. Because MAPK signaling pathways are involved in both pro- and anti-inflammatory pathways [54,55], therefore, we hypothesize that the observed response is facilitated by the activation of MAPK signaling pathways as indicated by an increased expression of genes in anti-inflammatory factors such as *Dusp1, Dusp5, Dusp6, Dusp10, Il1b*, and *Il1r2* that were unique to the IU region after sperm exposure. DUSP proteins are shown to be crucial for controlling inflammation and antimicrobial immune response [55]. Most strikingly, the majority of these genes were downregulated at 1.5 and 2.5 dpc when the embryos are present in the IU region, suggesting that sperm may play a role in the immunosuppression of the oviduct for safe embryonic development. However, this finding is in conflict with prior studies in the *in vitro* bovine oviductal epithelial cell (BOEC) culture model, in which sperm bind and induce transcripts involved in anti-inflammatory cytokines, such as *TGFB1* (transforming growth factor β1) and *IL10*, while decreasing proinflammatory transcripts such as *TNFa* (tumor necrosis factor α) and *IL1B* in the BOECs [56,57]. It is possible that this discrepancy could be due to differences between 1) *in vivo* vs. *in vitro* models or 2) murine vs. bovine model organisms. Our observed switch between activation and suppression of proinflammatory responses (or a switch from pro-to anti-inflammation) across the preimplantation period does not occur in pseudopregnant replicates, indicating that this phenomenon is a sperm- and embryo-mediated interaction. Most importantly, we established here, for the first time, that this observed pro- and anti-inflammatory response may be facilitated almost exclusively by the secretory cell population in the IU region when compared to their ciliated cell counterparts.

At 1.5 dpc, 2-cell embryos were located in the isthmus region. All embryos at later developmental stages1.5-2.5 dpc were stalled in lower isthmus and subsequently the UTJ region. At 3.5 dpc, almost all embryos transit the oviduct to the uterus. These findings are in agreement with previous findings in the mouse model [53,58]. Nutrients such as pyruvate, lactate, and amino acids are present in the oviductal fluid in several mammalian species [59-61]. After fertilization, 1-cell embryos acquire pyruvate and lactate for their energy source [62]. Then, the metabolism profile shifts from oxidative to glycolytic metabolism at later stages of preimplantation development [63,64]. Here we found that upregulated DEGs at 0.5 dpc were enriched for several energy metabolism biological processes, including pyruvate metabolic, glucose catabolic process to pyruvate, canonical glycolysis, and glycolytic process through glucose-6-phosphose. These pathways are subsequently downregulated between 1.5-3.5 dpc. We showed that genes involved in these pathways were unique to the IU region. Therefore, it is possible that the IU region is priming its environment to adjust for the production of specific energy sources required for early and late embryo metabolism as the embryo switches from utilizing pyruvate to utilizing glucose during successive developmental periods in the oviduct.

Lastly, we found that the oviduct was immunodynamic as indicated by the presence of immune cells as indicated by several immune markers such as neutrophils (*Ly6g*+), leukocyte (*Ptrpc*+), T cells (*Cd3d+, Cd3g+*), NK cells (*Nkg7+, Klrb1c+*), among others. This finding is in agreement with previous studies in human Fallopian tubes, as well as from our lab and other laboratories [15,65,66]. It suggests that the oviduct is capable of mounting innate immune responses in the presence of foreign bodies such as sperm.

In conclusion, we have demonstrated that the transcriptomic landscape of the oviduct at 4 different preimplantation timepoints was dynamic for both natural fertilization and pseudopregnancy using two independent cell/tissue isolations and sequencing techniques. Most novel findings from this study suggest that: 1) sperm were likely the key mediators in modulating inflammatory responses in the oviduct, potentially priming the oviduct to become tolerable to the presence of embryos, 2) inflammatory cytokine-mediated signals observed were likely generated by the secretory epithelial cells of the oviduct, 3) the oviduct is an immuno-dynamic organ, and 4) the oviduct could provide necessary nutrient enrichment in the luminal fluid at different stages of embryonic development. Currently, we are validating these findings at the translational levels using immunoblotting and cytokine arrays. In addition, we aim to determine luminal fluid contents including proteomic and other molecules that could be distinctly present at different stages of preimplantation period. Overall, our findings reveal an adaptive oviduct with unique transcriptomic profiles in different oviductal regions that may be specialized to influence sperm migration, fertilization, embryo transport and development. These findings could facilitate developments to ensure a proper microenvironment for the embryo to properly develop at each stage in an *in vitro* setting is established at the agricultural/clinical level.

## ACKNOWLEDGEMENT

The authors thank Kalli Stephens for helping maintain the C57BL/6J mouse colony and Gerardo Herrera for initial analysis of scRNA-seq data.

## AUTHORS’ CONTRIBUTIONS

R.M.F and W.W. designed the experiments. R.M.F. and D.J.C. performed the experiments. R.M.F., D.J.C., and W.W. analyzed the data and wrote, edited, reviewed and approved the final version of the manuscript.

